# Olfactory detection and discrimination in domestic dogs (*Canis lupus familiaris*)

**DOI:** 10.1101/2022.02.04.479113

**Authors:** Elodie Ferrando, Christoph D. Dahl

## Abstract

The extraordinary olfactory capabilities in detection and rescue dogs are well-known. However, the olfactory performance varies by breed and search environment (Jezierski et al., 2014), as well as by the quantity of training (Horowitz et al., 2013). While detection of an olfactory cue inherently demands a judgment regarding the presence or absence of a cue at a given location, olfactory discrimination requires an assessment of quantity, a task demanding more attention and, hence, decreasing reliability as an informational source (Horowitz et al., 2013). This study aims at gaining more clarity on detection and discrimination of olfactory cues in untrained dogs and in a variety of dog breeds. Using a two-alternative forced choice (2AFC) paradigm, we assessed olfactory detection scores by presenting a varied quantity of food reward under one or the other hidden cup, and discrimination scores by presenting two varied quantities of food reward under both hidden cups. We found relatively reliable detection performances across all breeds and limited discrimination abilities, modulated by breed. We discuss our findings in relation to the cognitive demands imposed by the tasks and the cephalic index of the dog breeds.

## Introduction

The dog’s high sensitivity of the olfactory system might subserve a particular fitness-enhancing trait present in the evolutionary ancestor of today’s canids, i.e. the detection of a prey in relative close proximity (Horowitz et al., 2013) or the inference about the location of a distal prey via airborne olfactory particles (Hepper and Wells, 2005). Apart from that, domestic dogs use olfaction in social contexts, such as social interactions (Bräuer and Blasi, 2021), identity recognition (Cafazzo et al., 2012; Lisberg and Snowdon, 2009), intrasexual competition for mates, territorial defence (Bidder et al., 2020) and parental care (Foyer et al., 2016; Lévy et al., 2004). In general, the olfactory bulb and its projection structures, including the olfactory tract and striae, are larger in dogs than in humans in absolute terms and relative to brain size (Kavoi and Jameela, 2011), suggesting higher olfactory functionality (Haehner et al., 2008). In numbers, olfactory receptors are 30 times more in dogs (*≈* 200 million) compared to humans (*≈* 6 million) (Horowitz, 2014; Lindsay, 2013). In particular, however, the actual size of the olfactory epithelium varies largely across breeds (Quignon et al., 2003), paralled by inter-breed differences in behavioural outcome: Dog breeds selected for olfactory tasks (scenting breeds) and wolves performed better in detecting one of four pots baited with a food reward than short-nosed and non-scent breeds (Polgár et al., 2016).

Despite a general consensus on the importance of olfaction in dog cognition, relatively little investigation has been conducted regarding some of the most fundamental perceptual abilities, such as discrimination of two scents, one of them stronger than the other (Becker et al., 1962; Cablk et al., 2008). Research programmes focused predominantly on visuo-cognitive questions, such as visual human-dog communication (Horowitz, 2021; Kerepesi et al., 2005) and interpretation of pointing gestures (Soproni et al., 2002). Likewise, most insights into quantity discrimination in dogs rose from *visual* numerical discrimination (Petrazzini and Wynne, 2016, 2017; Prato-Previde et al., 2008). These studies showed that dogs, including puppies (Petrazzini et al., 2020), tend to select significantly more often the visually presented large food quantity than the small food quantity in a choice task (Petrazzini and Wynne, 2016, 2017) or the visually presented larger amount of non-food items on two magnetic boards (Macpherson and Roberts, 2013), particularly for quanities of lower ratio (i.e. a large difference between the two quantities). This result conforms with Weber’s law (Fechner, 1860), where two comparable quantities (i.e. when the small quantity divided by the large quantity produces a large value) are harder to discriminate than two different quantities (i.e. when the small quantity divided by the large quantity produces a small value). This ratio effect has also been found in chimpanzees (Beran, 2001) and pigeons (Roberts, 2010) during visual quantity discrimination and is suggestive for an approximate number system (ANS) (Feigenson et al., 2004). The ANS is a cognitive representation system of numbers larger than four, allowing a spontaneous approximation of number of objects (Piazza, 2011) without relying on language or symbolic representations. The Weber fraction in the ratio effect is the just-noticeable difference in stimulus intensity and can be derived from the accuracy performance as the difference between the two closest discriminable numerosities (Piazza, 2011).The Weber fraction is relatively constant and a fixed proportion of the reference sensory level. It can be described as follows, where *I* is the original intensity of the stimulus and Δ*I* is the additional intensity to generate a noticeable difference (Equation 1):

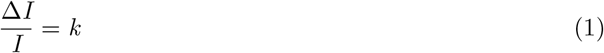

Due to the constancy of the Weber fraction, numerical differences at higher ratio (i.e. when the small quantity divided by the large quantity produces a large value) are typically perceived with lower accuracy. This generally accounts for the reported decline in discrimination performance with decreasing difference between quantities (Petrazzini and Wynne, 2016, 2017) and, in particular, for the reliance on the total amount rather than on the number for discriminating quantities (Petrazzini and Wynne, 2016). However, this interpretation is inconsistent with the view in the literature (Agrillo and Beran, 2013) that for numbers smaller than four a different numerical system than the ANS is engaged in the numerical representation, that is the object tracking system (OTS), wherein each perceived object is represented as a symbol, as a distinct unit, that adheres to the spatio-temporal principles of cohesion (Piazza, 2011): That is, such unit moves as a bounded whole and on connected unobstructed paths, principles that are conceptually related to the Gestalt laws in psychology (Ginsburg and Nicholls, 1988). Critically, the OTS is not affected by numerical ratio, i.e. 1:4 units can be discriminated with similar accuracy as 3:4 units. Therefore, the two-system explanation leads to a discrepancy between predicted and empirical outcomes, not only dog visual discrimination of quantities (Petrazzini and Wynne, 2016, 2017; Prato-Previde et al., 2008) but also in, for example, sea lions (Abramson et al., 2011), chimpanzees (Beran et al., 2008), and crows (Bogale et al., 2014). In response to that, a growing number of authors argue in favour of a single approximate number system that accounts for the complete number stream (Aulet et al., 2019; Beran, 2007; Beran and Parrish, 2016; Cantlon and Brannon, 2006; Cordes and Brannon, 2009).

Fewer studies addressed dogs’ capabilities of olfactorily discriminating quantities: On average dogs paid more attention to a covered plate baited with more food (5 units) than one baited with less food (1 unit), but did not select that plate significantly more frequently (Horowitz et al., 2013). In contrast, in a more recent study dogs were able to reliably discriminate five hot dog slices from one hot dog slice (Jackson, 2020; Jackson et al., 2021). In a separate experiment, quantity comparisons with varying food ratios showed that dogs did not adhere to the ratio effect, i.e. with increasing food ratio (e.g. 4:8 (low ratio) vs 4:6 (high ratio)) performance did *not* deteriorate, nor did they adhere to the distance effect, i.e. with decreasing difference in number of food units performance did *not* deteriorate (e.g. 4:8 (large difference) vs 2:4 (small difference)). Jackson (Jackson, 2020) suggested a difference in the processing of quantity discrimination between olfactory and visual domains. The lack of consistency in the assessment of dog’s capabilities of olfactory discrimination might, however, also suggest that olfactory capabilities in dogs are subjected to large inter-breed variability, where a simplified general model of dog olfaction does not apply rather than a breed-specific one: For some breeds with greater olfactory sensitivity, ratio differences (e.g. 1:5 vs 3:5) might well be perceived and proportionally well differentiated, whereas breeds with lower sensitivity might hardly be capable of telling apart the largest of a difference between quantities and would be indifferent to any modulation in food ratio.

In order to gain more consensus on dogs’ capabilities in olfactory discrimination of quantities and whether discrimination of quantities in the olfactory domain adheres to similar principles as the visual domain, we re-assess discrimination capabilities in a two-alternative forced choice (2AFC) paradigm. We also assess the detection rates of an olfactory food source, when the alternative was none in a 2AFC task. We hypothesise that dogs are better in detecting a food source than discriminating a larger from a smaller food source (*Hypothesis 1*). In this context, the detection task not only serves the purpose of replication of previous findings (Polgár et al., 2016), but it offers a means by which we can individually contrast detection scores of subjects and breeds with discrimination scores. Further, we hypothesise that dog olfactory capabilities vary largely between breeds, given the variability between breeds in a natural detection task (Polgár et al., 2016) and the olfactory receptor sequence polymorphism between breeds (Tacher et al., 2005) (*Hypothesis 2*). More specifically we predict a correlation of performance scores with the skull form (dolicho-to brachycephalic) of the dog breeds, following the general assumption that those breeds with a brachycephalic skull (i.e. the length of the craniumis shorter than the width) are worse in olfactory detection than breeds with mesocephalic or dolichocephalic skulls (Polgár et al., 2016) (*Hypothesis 3*). We also hypothesise that Weber’s law applies to dogs’ olfactory discrimination in general; however, we expect that some but not all breeds exhibit sufficient sensitivity to discriminate larger differences between quantities better than smaller ones (*Hypothesis 4*). For this reason, we chose a rather conservative experimental approach, where plates baited with food rewards were completely covered by a cup.

## Materials and methods

### Subjects

We used 41 domestic dogs (*Canis lupus familiaris*) (age in month: *µ* = 21.13, *σ* = 28.22) of the following breeds: Miniature Australian Shepherd (MAS), White Swiss Shepherd (WS), Jack Russel Terrier (JR), Siberian Husky (SH), Spitz (SP), Golden Retriever (GR), Bichon Bolognese (BB), French Bulldog (FB) (Table 1). All dogs were owned and raised by dog shelters in Switzerland and France. The dogs were naïve to the tasks and scientific procedures. Dogs were not fed within three hours prior to the experiment.

**Table 1:**
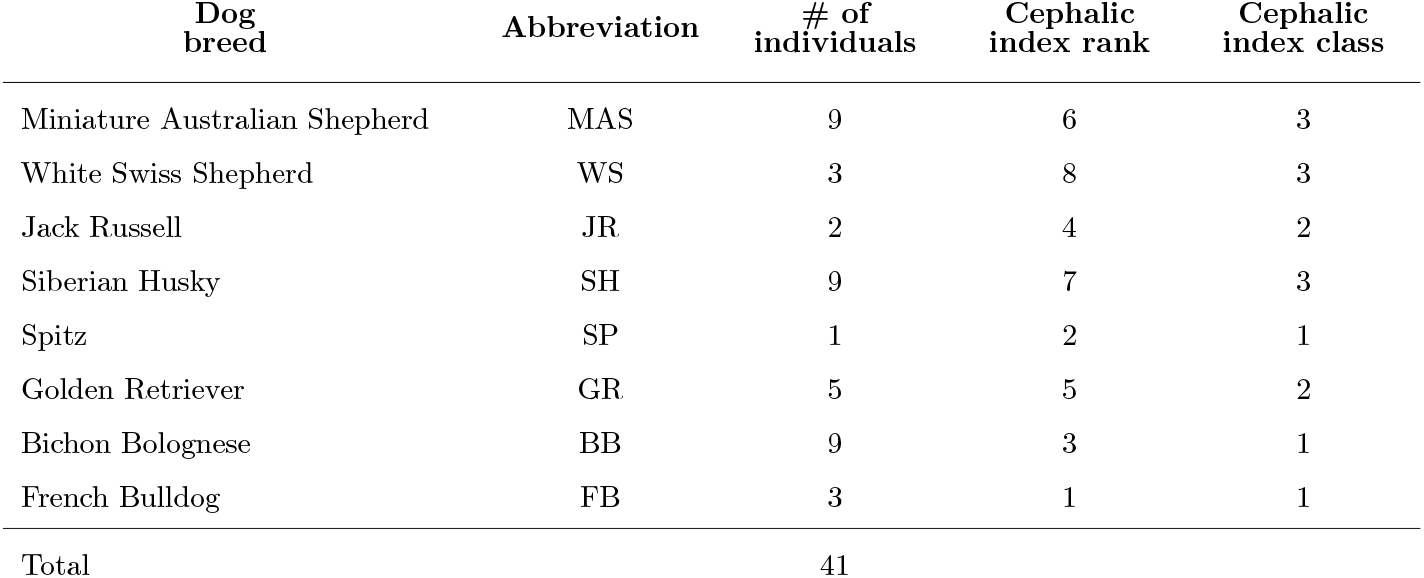
Dog breeds used in the study and assigned cephalic index rank (from ‘towards brachycephalic’ (rank 1) to ‘towards dolichocephalic’ (rank 8)) and class (from ‘towards brachycephalic’ (class 1) to ‘towards dolichocephalic’ (class 3)).

### Apparatus

All dogs were tested in a comparable fashion in enclosed and sheltered outdoor testing arena, measuring approximately 20 - 40 m^2^. Using natural ventilation, where two sides of the testing arena were partially opened, we ensured that the testing arena was not saturated with odours. For the experiments the dogs were separated from other dogs and had no visual or acoustic contact. Water was always available during the experiment. We used glass plates (100 by 100mm base plate, 3mm thick, with little knobs to keep the plate 3mm above ground) and plastic cups (cup opening diameter: 67mm; lower diameter: 45mm; height: 100mm; thickness: .5mm) for a controlled stimulus presentation (see Procedure).

### Procedure

Dogs were tested in two tasks: (1) *detection* task, (2) *discrimination* task. In both of these tasks, we used a two-alternative forced choice (2AFC) paradigm, where (1) a food reward of varying quantity was placed below one of two cups at 1 m distance, requiring the dog to indicate which of the cups contains the bait by approaching the respective cup, or (2) two food rewards of varying quantities and reward ratios were placed below both cups at 1 m distance, requiring the dog to indicate its preference by approaching the cup, assuming it would naturally choose the cup with the strongest odour and therfore the larger quantity of food reward.

Prior to the testing procedure, dogs underwent a habituation phase, where they were familiarised with items used in the experiments, i.e. cups and glass plates, and where they could freely explore the testing environment. They were also habituated to the human experimenters. After that, each dog was trained on the general task requirements by running four *detection* trials, where a food unit was placed under one of two cups and the dog was required to indicate which of the two cups it wants to get uncovered by approaching the cup of its choice. The operational definition of a choice was as follows: The dog needs to be in an area of 30cm or closer to the cup, remain there for 3 seconds or touch the cup.

In the testing phase, all dogs did both, the *detection* and the *discrimination* tasks. In both tasks, the dog’s starting point was outside of the testing arena, kept on a leash by Experimenter 1. Experimenter 2 turned two cups upside-down onto two plates and placed a food reward under one of the cups (task 1) or two food rewards under both cups (task 2) according to the experimental protocol. Thereby, the food reward(s) were fully covered by the cup(s). The dog, guided by Experimenter 1, entered the arena at the other end and was brought to the starting point of the trial, which was at a distance of 1.5 m to the midpoint of the cup arrangement. The dog was aligned equidistantly to the cups, forming a perpendicular bisector to the cup alignment. In the beginning of each trial, the dog was held back for 10 sec, allowing it to sense the reward location at the starting point, and then released by Experimenter 1 to freely approach the target location. The dog then indicated its choice, and Experimenter 2 lifted up both cups simultaneously. If the dog’s choice was correct, the dog was allowed to eat the food reward under the cup chosen. If the dog’s choice was incorrect, the dog did not receive a reward. Experimenter 1 then led the dog out of the arena. Experimenter 2 replaced the cups with new and unused cups and the glass plates with clean glass plates, and cleaned the environment, if necessary.

The reward-position was counter-balanced in the *detection* task and the larger reward-position was counter-balanced in the *discrimination* task. In the *detection* task, 1, 2 or 3 food reward units were used; in the *discrimination* task the following food reward units were compared: 1:2; 1:3; 1:4. Each reward unit was a standardised half-slice of Cervelat sausage, a typical Swiss sausage containing equal parts of beef, pork, bacon, and pork rind, along with spices, curing salt and cutter additives.

We further ensure that there was no experimenter-bias by having Experimenter 2, which controlled the cup placement and the uncovering of the reward, looking down at the floor during the trial. Experimenter 1 was blind regarding the positioning of the food reward. In the *detection* task, each dog did three sessions on three different days, each containing 2 trials per quantity of food reward, hence 6 trials per session and a total of 18 trials per dog. In the *discrimination* task, each dog did three sessions on three different days, each containing 2 trials per food reward ratio (1:2, 1:3, 1:4), hence 6 trials per session and a total of 18 trials per dog. The trials were randomised for each dog. All experimental sessions were digitally recorded using two Panasonic^®^ HC-V160 cameras (Panasonic^®^ Corporation, Kadoma, Osaka, Japan).

### Data analysis

Each trial resulted in either a correct detection (*detection* task), where correct was coded as ‘1’ and incorrect as ‘0’, or in a ‘correct’ discrimination (*discrimination* task), where ‘1’ indicates large quantity of food reward and ‘0’ small quantity of food reward. We are aware that for the discrimination task there is no ‘correct’ or ‘incorrect’ response. However, for simplicity, we refer to the responses as ‘correct’ or ‘incorrect’, assuming that dogs’ natural choice would aim at maximising the quantity of food reward, hence they would prefer the larger quantity in the discrimination task over the smaller quantity (Ward and Smuts, 2007). Any outcome from the *discrimination* task needs to be discussed under this assumption. Henceforth, we refer to the response variable as *performance* (correct vs incorrect).

Analyses were performed in Matlab (Mathworks^®^, Natick, Massachusetts, USA). We fitted a generalised linear mixed-effects model (fitglme function in Matlab) with binomial error structure and logit link function to our complete dataset (*detection* and *discrimination* task). We followed a model fitting procedure commonly accepted in the literature (Dobson and Barnett, 2018; Fedurek et al., 2019). To fully account for *performance* as response variable, we fitted four predictor variables: (1) *breed* of dog, (2) *quantity* of reward, (3) *side* of presentation, and (4) *task* (*detection* vs *discrimination*). Since we were interested in differentiating those effects that are similar across breeds from those that are different between breeds, we fitted the two-way interactions between *breed* and the remaining three main predictors. We also fitted the remaining interactions between factors *task, side*, and *quantity*. We assumed that there might be performance differences between breeds regarding discrimination of *quantity* of reward depending on the *task*. We also assumed that there might be a *side* response bias that depends on (1) *breed* and *task*, (2) *breed* and *quantity*, and (3) *quantity* and *task*. A side bias is a stereotypical choice behaviour whose prevalence can be increased under conditions of lower stimulus discriminability (Trevino, 2014). The rationale for assuming a side bias is that, if certain dog breeds were less likely to discriminate two given olfactory cues, these breeds might be more likely to develop a side bias. Hence, we fitted four additional three-way interactions. Thus, the full model contained six two-way interactions and four three-way interactions. In the next step, we fitted (1) random intercepts to *subject*, and random slopes and intercepts to the terms (2) *side* of presentation in *subject*, (3) *task* in *subject* and (4) *quantity* in *subject*, since we predicted that (1) individual dogs perform overall differently, (2) are potentially subject to a side bias, (3) perform better in one task than the other, and (4) some individuals’ performance might depend on the presented quantity.

In a second model, we fitted four predictor variables, replacing *breed* with *cephalic index* (CI), since we predicted that dogs’s performance correlates with their skull form (dolicho-to brachycephalic). To this end, we determined the cephalic index for each breed according to the literature (Evans and De Lahunta, 2013; McGreevy et al., 2013; Stone et al., 2016; Teng et al., 2016), reconfirmed the order of breeds with regard to the cephalic index with an expert (Zśofia Bognár, personal communication) at the Department of Ethology of ELTE Eötvös Loránd University, ranked the breeds regarding the resulting cephalic index (‘Cephalic index rank’, Table 1) and binned them into three classes (‘Cephalic index class’, Table 1). We do not assign commonly known labels, such as brachy-, meso-, or dolichocephalic to individual breeds, rather than relative class labels, such as ‘*towards* brachycephalic’ (class 1), or ‘*towards* dolichocephalic’ (class 3). More information on the model fitting procedure can be found in the supplementary materials. In addition, we used Pearson correlation to determine the extent to which subjects performance scores in the *detection* and *discrimination* task were correlated.

## Results

### Descriptive analysis

Figure 1A-D shows the proportion correct responses over all dogs and breeds combined (A) and for each breed separately (B) in the *detection* task. Overall, dogs were correct in 70 to 75% of their choices (Figure 1A). We found a considerable variation between breeds, with Jack Russell Terriers, Siberian Huskies, and French Bulldogs showing relatively high performance scores and Bichon Bolognese showing relatively low performance scores that even fell below the chance level of 50% correct detection (Figure 1B). It needs to be noted, however, that some breeds were represented by a low number of subjects. Factoring in the quantity of food reward, we found an overall increase of performance with increasing quantity of food (Figure 1C). At the level of individual breeds, however, there was little systematicity in terms of the predicted performance increase with increasing quantity of food reward (Figure 1D): The only breed that followed the prediction was the Siberian Husky.

**Figure caption 1:**
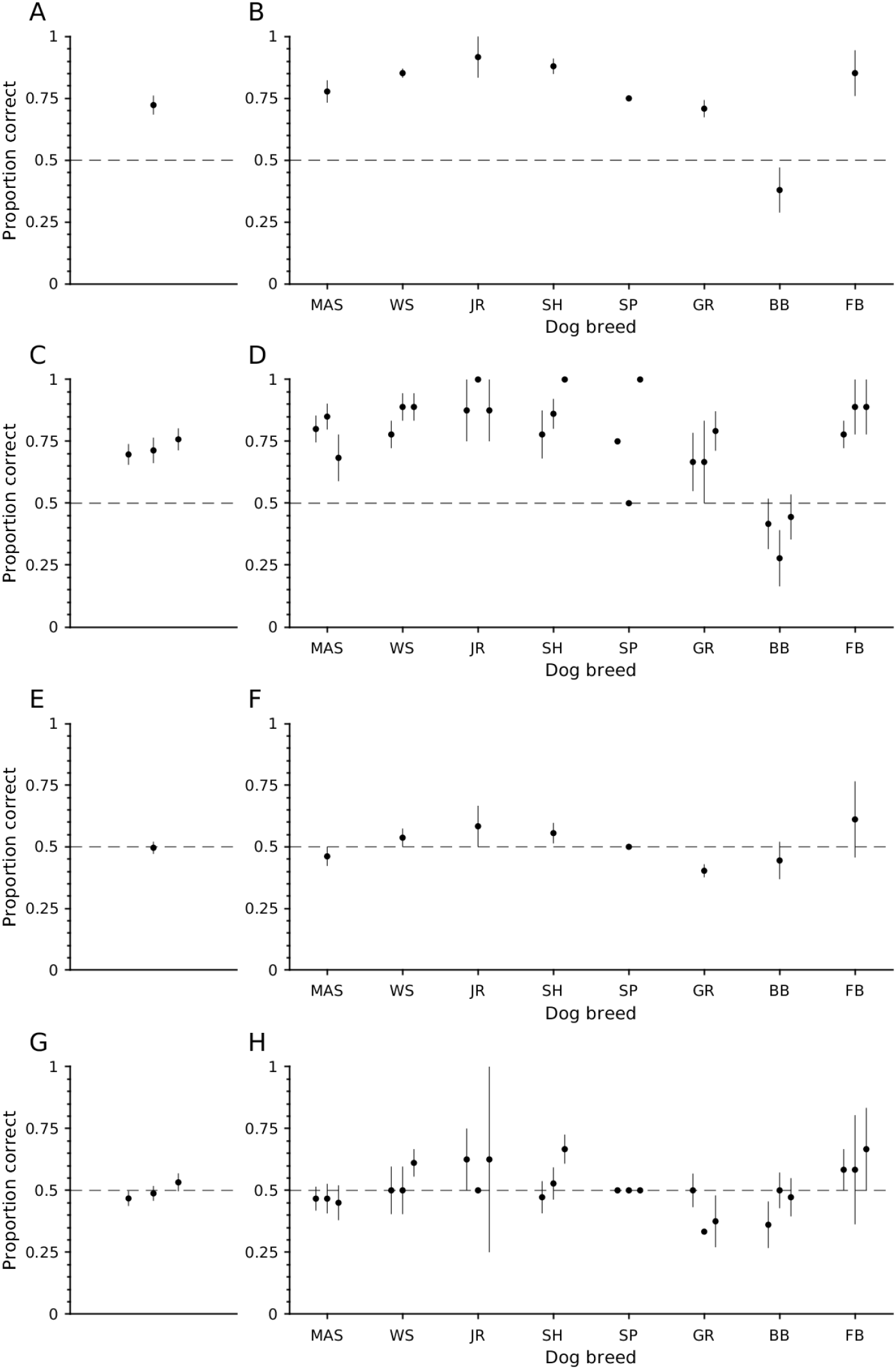
Detection and discrimination tasks. All panels show the proportion correct responses in the detection task on the y-axis. The black dots represent the mean and the whiskers the standard error of the mean. A-D. Detection task. A shows the overall performance (grand mean); B shows the performances for different breeds (x-axis); C and D are similar to A and B, but show the performance as a function of quantity of food reward (C) for each breed (D). For each set of three means and standard errors of the mean, the smallest food reward (1 unit) is shown to the left and the largest (3 units) to the right. E-H. Discrimination task. E shows the overall performance (grand mean); F shows the performances for different breeds (x-axis); G and H are similar to E and F, but show the performance as a function of quantity of food reward (G) for each breed (H).

Figure 1E-H shows the proportion correct responses over all dogs and breeds combined (E) and for each breed separately (F) in the *discrimination* task. Overall, dogs were correct in 50% of their choices, thus, at chance level (Figure 1E). There was variation between breeds, with the same breeds performing marginally better as in the *detection* task (i.e., Jack Russell Terriers, Siberian Huskies, and French Bulldogs, while the latter showed large within-breed variation (Figure 1F). There were some breeds that performed below chance level, i.e. Miniature Australian Shepherd, Golden Retriever, and Bichon Bolognese. All the differences in the *discrimination* task were marginal. The choice accuracy varied as a function of quantity of food reward, reflected in a small performance increase with increasing difference between the smaller and the larger quantities at the grand mean level (Figure 1G). Also at the level of breeds, some breeds showed a systematic increase of performance with increasing difference between the smaller and the larger quantities: The breeds that followed the prediction most closely were the Siberian Husky and the French Bulldog (Figure 1H).

### Model investigation

In the *breed model*, we found that *breed* s performed differently (LRT: 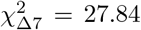, *p <* 0.001; breed model: df = 55, AIC = 1438.3), and the two *task* s yielded different overall results (LRT: 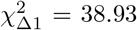, *p <* 0.001; task model: df = 61, AIC = 1461.4). These effects were not independent of each other, resulting in a significant interaction of *breed* and *task* (LRT: 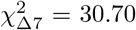 *p <* 0.001; breed-task model: df = 55, AIC = 1441.1). Furthermore, we found a trend regarding the factor *quantity* (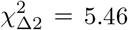, *p =* 0.007; quantity model: df = 60, AIC = 1425.9). The interaction between factors *breed* and *quantity* was not significant, however close to a trend (LRT: 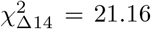, *p =* 0.10; breed-quantity model: df = 48, AIC = 1417.6). The factor *side* and the remaining interactions were non-significant (Supplementary materials, Supplementary table 1). Random effects revealed that subjects performed at equal levels regarding the *task* involved and the *quantity* of food reward tested on: There was no significant interaction between the intercept *subject* and the slope grouped by *task* (LRT: 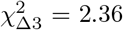, *p =* 0.50; subject-task model: df = 58, AIC = 1418.8), nor was there a significant interaction between the intercept *subject* and the slope grouped by *quantity* (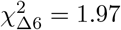,*p =* 0.92; subject-quantity model: df = 55, AIC = 1412.4). One Bichon was significantly different in its responses between the two *task* s (see ‘*Subject* x *Task* ‘ in Supplementary table 2). Some subjects, however, showed a side bias, reflected in a correlation between the intercept grouped by *subject* and the slope grouped by *side* (LRT: 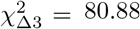, *p <* 0.001; subject-side model: df = 58, AIC = 1497.3; see ‘*Subject* x *Side*’ in Supplementary table 2). Significantly different subjects were 4 Miniature Australian Shepherds, 2 French Bulldogs, 5 Bichons, and 1 Siberian husky. Adding the random factor *subject* with its interactions *side* improved the model, and subject variability in this regard cannot be neglected. There was, however, a deterioration in modelling the interaction between the intercept *subject* and the slope *quantity* and *task* (see above) as opposed to omitting these factors from the model, suggesting that there was no systematic difference regarding the discrimination performance as a function of *quantity* or *task* at the *subject* level.

The second model, modelling the factor *cephalic index* and replacing the factor *breed*, showed a significant modulation by the factor *cephalic index* (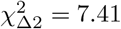, *p <* 0.05; cephalic index model: df = 30, AIC = 1413.1). In more detail, *cephalic index* class 1 was significantly different from *cephalic index* class 3 (Figure 2A; Supplementary table 3). As expected from the previous model, the factor *task* was significant (LRT: 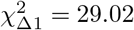, *p <* 0.001; task model: df = 31, AIC = 1436.7; Supplementary table 3) as well as the interaction of *cephalic index* and *task* (LRT: 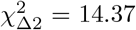, *p <* 0.001; task model: df = 30, AIC = 1420; Figure 2B; Supplementary table 3). This interaction manifested in an increase of performance scores with increasing *cephalic index* class, i.e. from brachycephalic to dolichocephalic, in the *discrimination* task, but remains at similar levels across *cephalic index* classes in the *discrimination* task. Remaining factors follow the same directions as in the first model based on the factor *breed*. The parameter estimates of the model based on the factor *cephalic index* are summarised in the Supplementary table 3.

**Figure caption 2:**
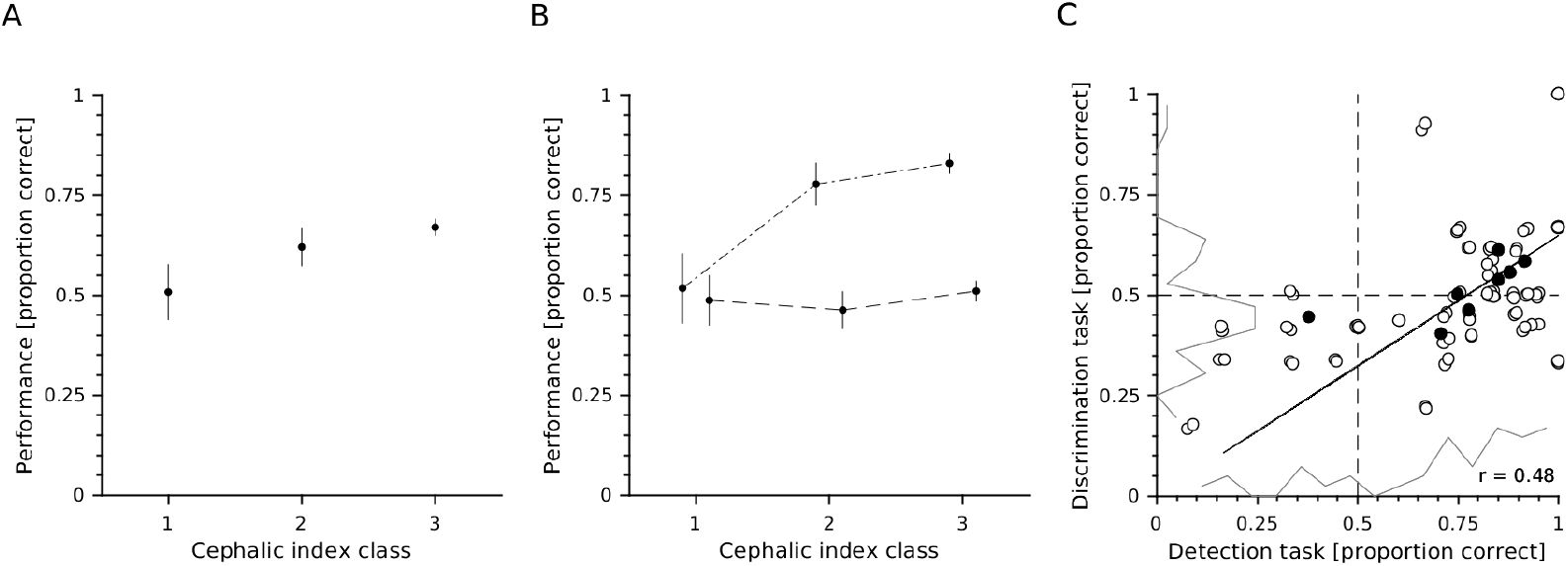
Cepahlic index classes, and correlation of detection and discrimination performance scores. A. The general perfomance scores are shown as a function of the cephalic index classes. Class labels on the x-axis stand for ‘towards brachycephalic’ (class 1) and ‘towards dolichocephalic’ (class 3). Shown are the means and standard errors of the mean across individuals. B. Similar as in A, but for the *detection* task (dash-dotted line) *discrimination* task (dashed line), respectively. C. Open circles indicate individual subjects’ performance in the *detection* task (x-axis) and the *discrimination* task (y-axis). The regression line (solid black line) was calculated on the basis of the individual subjects’ performance scores. For illustration, filled black circles indicate the mean across subjects for each breed. The solid grey lines show the distribution of data samples, computed via histograms containing 20 equally large bins. The dashed black lines indicate the 50% chance level for both tasks.

We further found that the *detection* and *discrimination* tasks correlated at the subject level (*r* (39) = 0.48, *p <* 0.01; Figure 2C).

## Discussion

In two experiments we studied the olfactory capabilities of untrained domestic dogs of different breeds, using a two-alternative forced choice (2AFC) paradigm. We hypothesised that, in general, dogs perform better in the detection of an olfactory cue, i.e. when no cue is the alternative choice, than in the discrimination of a larger olfactory cue from a smaller one (*Hypothesis 1*). Further, we hypothesised that given the genetic diversity of the canine olfactory system (Robin et al., 2009), dog breeds vary in olfactory capability, similar to a naturally occuring variance in visual capacity between breeds (McGreevy et al., 2004) (*Hypothesis 2*). We also predicted an influence on performance scores in both tasks depending on the skull form of the dog breed, i.e. dolicho-, meso-or brachycephalic, as a result of selective breeding causing functional anatomical changes in the brain (*Hypothesis 3*). Lastly, we also hypothesised that Weber’s law applies to dogs’ olfactory discrimination in general; however, due to our relatively conservative methodological approach, i.e. fully cover the food reward, some but not all breeds exhibit sufficient sensitivity to discriminate larger differences between quantities better than smaller ones, and hence, not all breeds might adhere to Weber’s law under the given experimental circumstances (*Hypothesis 4*).

Addressing *Hypothesis 1*, we found that the detection and discrimination tasks produced largely different outcomes: While most breeds performed relatively well in detecting an olfactory stimulus, the discrimination of two olfactory stimuli varying in intensity (one being a larger quantity of food reward than the other) yielded about 50% correct responses - paralleling performance at chance level. This performance difference comes at no surpise, given the differences in computational demands between *detection* and *discrimination*. While in the *discrimination* task, subjects are required to discriminate the qualities of stimuli identities, i.e. the magnitudes of two smells, in the *detection* task, subjects are required to simply detect the presence of an identity, the smell, irrespective of the quality of the identity, i.e. the magnitude of smell (Schópper et al., 2020). The relatively low discrimination performance is in accordance with Horowitz and colleagues (Horowitz et al., 2013), who found that dogs chose the larger quantity of food reward more often (61%), but not significantly more than the smaller quantity. Dogs in their study were given the chance to explore two covered plates for about 3 seconds each, either in sequential or in simultaenous presentation. This procedure is different from ours, where dogs were relying on sniffing the larger quantity from 1 m distance and then indicating the cup of choice by approaching it. Our methodological approach, and, in particular, fully covering the food reward(s), resulted in higher task difficulty, given our intention to amplify and map out inter-breed differences. This more conservative approach might, thus, explain the even lower average performance scores during the *detection* and *discrimination* tasks in our study than in Polgár et al. (2016) and Horowitz et al. (2013), respectively. Notably, the rather low *olfactory* capability in our study and the study by Horowitz et al. (2013) is in contrast to Jackson and colleagues’ study (Jackson et al., 2021), that showed successful olfactory discrimination, but failed to show that dogs adhere to the ratio and/or distance effect. Critically, in their study, the containers had four parallel slits, allowing air to flow through the base of the container and olfactory particle flux. Our study and the study by Horowitz et al. (2013) is also in contrast to dogs’ *visual* capability in discriminating two food quantities (Petrazzini and Wynne, 2016; Prato-Previde et al., 2008; Ward and Smuts, 2007). This contrast between sensory modalities does not necessarily suggest that dogs discriminate quantities predominantly relying on vision or that neural mechanisms for olfactory discrimination do no exist, rather than that the visual and the olfactory tasks are by no means comparable with regard to the necessary magnitude of stimulation to generate a just-noticeable-difference. In other words, the Weber fraction varies between sensory modalities. Along the same lines, it might be that in a more naturalistic scenario where visual and olfactory stimulation co-occur the information from both modalities converge into a multisensory representation, potentially leading to multisensory enhancement, generating a more coherent representation based on which a more well-informed decision can be taken (Franzen et al., 2020; Mercier and Cappe, 2020). This redundancy principle, i.e. acquiring overlapping information from the environment, is a well-known design principles of intelligent systems and leads to systemic robustness and behavioural diversity (Pfeifer et al., 2005). In other words, the decision is made on the basis of multisensory rather than unisensory representations, leaving unisensory testing methods with little ecological relevance. This said, an explanation for why visual discrimination appears to be more sensitive than olfactory discrimination might be that the two modalities are complementary in the sense that, while the olfactory domain provides directional information of a (larger) prey in a very coarse manner, the visual domain provides the means to spot the (larger) prey at very high spatial and temporal resolution. Hence, there is no evolutionary pressure for an olfactory system with higher spatial resolution given the high precision of the visual system. Another explanation, which is not mutually exclusive to the above, is that dog olfaction does not comply with the notion of two systems, i.e. the ANS and OTS, as suggested in the visual domain, but rather relies on an all-or-nothing response system (Jackson, 2020). While in most sensory modalities the underlying physical principles are well understood; in olfaction, it remains unknown what principles specific chemical compounds in the air, forming odours, adhere to (Pannunzi and Nowotny, 2019). Hence, the lack of knowledge regarding the physical principles of odours prevents us from prematurely concluding that the olfactory system underlies similar cognitive mechanisms as the visual system regarding numerosity. Also, the fact that, in our experiment and in Horowitz et al. (2013), the olfactory stimuli were covered during presentation further augments the inability to precisely control the odours experimentally and to infer physical properties like concentration and spatio-temporal distribution.

Addressing *Hypothesis 2*, we found that detection and discrimination performances also varied largely between breeds. In detail, Jack Russell Terriers performed best in the *detection* task, followed by French Bulldogs, Siberian Huskies and White Swiss Shepherd. French Bulldogs and Jack Russell Terriers also performed best in the *discrimination* task, followed closely by Siberian Huskies and White Swiss Shepherd. When directly comparing olfactory *detection* and *discrimination* performances, individual subjects were positively correlated, suggesting that those individuals that were better in the *detection* task were also better in the *discrimination* task. This supports the notion that, in theory, dogs possess the capability of discrimination of olfactorily presented quantities, however, in our experiment the requirements for the *discrimination* task were too high, i.e. the differences between two presented olfactory cues remained largely undetected due to fully covering the food rewards. This directly highlights the additional task demands, when synchronously evaluating two olfactory sources as opposed to localising one: While keeping the ratio constant between tasks (i.e. we presented single cues at magnitudes of 1, 2, and 3 in the *detection* and two cues at magnitudes of 1 and 2, 1 and 3, and 1 and 4 in the *discrimination* task), the performance scores dropped about 25% between the *detection* and *discrimination* tasks. It needs to be noted that we did not aim at highest performance scores rather than at detectable differences between the two tasks and breeds. Also, systematically adapting the sensitivity range, potentially leads to a decrease of the Weber ratio to a degree where mathematically a discrimination task approximates a detection task, i.e. when the smaller quantity is marginally small and fully overshadowed by the large quantity, and, hence, conceptually it becomes increasingly difficult to distinguish a *discrimination* task from a *detection* task. This is particularly problematic without a solid understanding regarding the spatio-temporal distribution of odours (Pannunzi and Nowotny, 2019).

Addressing *Hypothesis 3*, we found that dogs of different *cephalic index* classes performed differently overall as well as a function of the *task*. We are aware that our classification of dog breeds into three classes does not follow the scientifically accepted classification of breeds into brachy-, meso-and dolichocephalic types of breeds, but given the limited number of breeds used in our study and the relatively centralised, i.e. mesocephalic, distribution, we decided to proportionally scale them within the pool of breeds used in this study. Clearly, a more detailed assessment on how cephalic type influences the olfactory *detection* and *discrimination* performances is required. Nevertheless, our analyses revealed that dog breeds differed in their abilities to detect and discriminate olfactory cues not arbitrarily, but in accordance with the general assumption that those breeds with a brachycephalic skull (i.e. the length of the cranium is shorter than the width) are worse in olfactory detection than breeds with mesocephalic or dolichocephalic skulls (Polgár et al., 2016), but see (Hall et al., 2015). It has been argued that selective breeding for reduced faces not only resulted in functional anatomical changes in the nose, but in a reduction in brain matter devoted to olfaction and spatial reorganisation of the brain (Selba et al., 2021). While a reduction of olfactory cortical areas clearly impacts the associated functions, such as olfactory detection and discrimination, the impact of spatial reorganisation of the brain areas and associated functions are currently investigated: Anatomical differences between breeds as a result of selective breeding and reflected in the cepahlic index have been reported (Hecht et al., 2019; Selba et al., 2021).

Addressing *Hypothesis 4*, we found that statistically dogs did not adhere to the ratio effect, i.e. their discrimination performance did not improve with decreasing Weber ratio or increasing differences between the quantities presented. This finding is in accordance to Jackson et al. (2021), who also found no evidence for a ratio effect. Nevertheless, in our study, the factor *quantity* played a crucial role in the model fitting and cannot be ignored due to its *p*-value being close to significance (7%). Indidividual dog breeds, such as the Siberian Husky, indeed reflect this trend very clearly: While trials with small quantity differences yielded lower correct discrimination, those trials with larger quantity difference yielded higher correct discrimination. This finding in the olfactory domain parallels findings in the visual domain (Petrazzini and Wynne, 2016, 2017) and suggests similar underlying principles for the discrimination of two magnitudes in the visual and the olfactory sensory modalities. The finding supports the idea of an olfactory equivalent to the approximate number system (ANS) (Feigenson et al., 2004) in the visual domain, which due to the nature of olfactory stimulation most likely adheres to an approximation of olfactory intensity throughout the complete intensity scale, and is therefore in contrast to the idea of two number systems, the ANS for numbers larger than four and the object tracking system (OTS) for numbers smaller than four as proposed in the visual domain.

Furthermore, it turned out that *subject* as random factor does not improve the model-fitting. This suggests inter-individual performance differences do not systematically occur neither at the level of *task* -related performances nor in dependence on food *quantity*. However, a limited number of subjects showed *side*-biases that led to a significant interaction of *subject* and *side*. This side-bias can be interpreted as a coping strategy for increasingly difficult task requirements and might also reduce with less conservative experimental methods. Future studies may address performance differences in olfactory detection and discrimination with regard to differences in temperament in general and motivational traits in particular between breeds and across individuals.

## Acknowledgements

We are grateful for the Startup-funding of Taipei Medical University (108-6402-004-112) and the Taiwan Ministry of Science and Technology research grant (110-2311-B-038-002) awarded to CDD. We thank Guillaume Dezecache, Ph.D., for comments on the manuscript.

## Author contributions

EF: study design, data collection, acquisition of resources; CDD: study design, analysis and interpretation, writing article.

## Ethical approval

All applicable international, national, and/or institutional guidelines for the care and use of animals were followed, i.e., the Swiss law on animal protection and welfare. This study was approved by the Swiss Federal Veterinary Office (approval number VD3383) for experiments conducted in Switzerland. According to the local authorities (Comité d’Ethique de l’Expérimentation Animale Grand Campus Dijon, Université de Bourgogne, Maison de l’Université, Esplanade Erasme, 21078 Dijon, France), non-invasive studies on dogs are allowed to be conducted without any special permission in France.

