## Supplementary materials for "Olfactory detection and discrimination in domestic dogs (*Canis lupus familiaris*)"

#### An investigation on the olfactory capabilities of domestic dogs (*Canis lupus familiaris*)

### Supplementary materials

#### Materials and methods

For both models, the *breed* model and the *cephalic index* model, we created a null model, which consisted of the similar structure as the corresponding full model, however leaving only *breed* or *cephalic index*, respectively, as fixed effect, while preserving all random effects. Using likelihood ratio test (LRT), we compared the null model with the full model. Assuming a significant improvement for the full model over the null model, the non-significant interaction terms were removed from the full model, reaching a model containing only significant interaction terms and both significant and non-significant main effects (Hector et al., 2010; Forstmeier and Schielzeth, 2011), henceforth referred to as the final model. Evaluation of fixed effect were on the basis of the *final* model.

This procedure resulted in the following models (Wilkinson notation):

*Full breed model*:

'Response ∼ 1 + Breed + Quantity + Side + Task + Breed:Quantity + Breed:Side + Breed:Task + Side:Task + Quantity:Task + Quantity:Side + Breed:Side:Task + Breed:Quantity:Task + Breed:Quantity:Side + Quantity:Side:Task + (1|Subject) + (Side|Subject) + (Task|Subject) + (Quantity|Subject)'

*Null breed model*:

'Response ∼ 1 + Breed + (1|Subject) + (Side|Subject) + (Task|Subject) + (Quantity|Subject)'

*Final breed model*:

'Response ∼ 1 + Breed + Quantity + Side + Task + Breed:Quantity + Breed:Task + Quantity:Task + Breed:Quantity:Task + (1|Subject) + (Side|Subject) + (Task|Subject) + (Quantity|Subject)'

*Final cephalic index model*:

'Response ∼ 1 + Cephalic index + Quantity + Side + Task + Cephalic index:Quantity + Cephalic index:Task + Quantity:Task + Cephalic index:Quantity:Task + (1|Subject) + (Side|Subject) + (Task|Subject) + (Quantity|Subject)'

#### Results

The *full breed model*, consisting of four fixed factors and their interactions that may explain the dogs' *performance* in the two tasks, was significantly different from the *null breed model* (LRT: $\chi^2_{\Delta 74} = 154.51$, $p < 0.001$; null model: df = 20, AIC = 1436.6). In addition to the reported results in the main manuscript, we found the following non-significant results: The factor *side* was not significant (LRT: $\chi^2_{\Delta 1} = 2.51$, $p = 0.11$; side model: df = 61, AIC = 1424.9). The remainig interactions were non-significant (Two-way interaction *quantity* and *task*: LRT: $\chi^2_{\Delta 2} = 3.10$, $p = 0.21$; quantity-task model: df = 60, AIC = 1423.5; three-way interaction *breed*, *quantity* and *task*: LRT: $\chi^2_{\Delta 14} = 19.45$, $p = 0.15$; breed-quantity-task model: df = 48, AIC = 1415.9). The parameter estimates of the final model are summarised in the Supplementary table 1.

### Tables

*Supplementary table 1: Results of the model investigating performance scores in detection and discrimination tasks. The table contains parameter estimates for the final model based on the factor 'breed' (first model).*

| | | **Estimate** | **SE** | ***t*-stat** | **DF** | ***p*-value** | **CI (95%) [lower, upper]** |
|---|---|---|---|---|---|---|---|
| | Intercept | 0.48 | 0.10 | 4.88 | 1185 | 0.001 | [0.29 0.67] |
| **Breed** | Miniature Australian Shepherd (MAS) | -0.05 | 0.15 | -0.31 | 1185 | 0.76 | [-0.35 0.25] |
| | White Swiss Shepherd (WS) | 0.14 | 0.24 | 0.59 | 1185 | 0.55 | [-0.33 0.61] |
| | Jack Russell (JR) | 0.41 | 0.32 | 1.28 | 1185 | 0.20 | [-0.22 1.04] |
| | Siberian Husky (SH) | 0.35 | 0.17 | 2.03 | 1185 | 0.04 | [0.01 0.69] |
| | Spitz (SP) | -0.09 | 0.41 | -0.23 | 1185 | 0.82 | [-0.89 0.71] |
| | Golden Retriever (GR) | -0.30 | 0.21 | -1.47 | 1185 | 0.14 | [-0.71 0.10] |
| | Bichon Bolognese (BB) | -0.80 | 0.17 | -4.75 | 1185 | 0.001 | [-1.13 -0.47] |
| | (against French Bulldog (FB)) | | | | | | |
| **Quantity** | Quantity Δ1 (1 unit) | -0.10 | 0.09 | -1.19 | 1185 | 0.23 | [-0.27 0.07] |
| | Quantity Δ2 (2 units) | -0.07 | 0.08 | -0.84 | 1185 | 0.40 | [-0.23 0.09] |
| | (against Quantity Δ3 (3 units)) | | | | | | |
| **Side** | Right | -0.16 | 0.10 | -1.61 | 1185 | 0.11 | [-0.34 0.03] |
| | (against Left) | | | | | | |
| **Task** | Detection task | 0.43 | 0.06 | 6.92 | 1185 | 0.001 | [0.31 0.55] |
| | (against Discrimination task) | | | | | | |
| **Breed x Task** | Miniature Australian Shepherd (MAS) : Detection | 0.07 | 0.09 | 0.79 | 1185 | 0.43 | [-0.11 0.26] |
| | White Swiss Shepherd (WS) : Detection | 0.08 | 0.14 | 0.53 | 1185 | 0.60 | [-0.21 0.36] |
| | Jack Russell (JR) : Detection | 0.25 | 0.23 | 1.09 | 1185 | 0.28 | [-0.20 0.70] |
| | Siberian Husky (SH) : Detection | 0.22 | 0.12 | 1.79 | 1185 | 0.07 | [-0.02 0.45] |
| | Spitz (SP) : Detection | -0.06 | 0.26 | -0.22 | 1185 | 0.82 | [-0.57 0.45] |
| | Golden Retriever (GR) : Detection | -0.02 | 0.12 | -0.13 | 1185 | 0.90 | [-0.25 0.22] |
| | Bichon Bolognese (BB): Detection | -0.55 | 0.11 | -4.99 | 1185 | 0.001 | [-0.77 -0.33] |
| | (against French Bulldog (FB) and Discrimination task) | | | | | | |
| **Breed x Quantity** | Miniature Australian Shepherd (MAS) : Quantity Δ1 (1 unit) | 0.14 | 0.13 | 1.08 | 1185 | 0.28 | [-0.11 0.40] |
| | White Swiss Shepherd (WS) : Quantity Δ1 (1 unit) | -0.10 | 0.20 | -0.46 | 1185 | 0.64 | [-0.48 0.30] |
| | Jack Russell (JR) : Quantity Δ1 (1 unit) | 0.10 | 0.30 | 0.34 | 1185 | 0.74 | [-0.49 0.69] |
| | Siberian Husky (SH) : Quantity Δ1 (1 unit) | -0.31 | 0.16 | -1.93 | 1185 | 0.05 | [-0.62 0.01] |
| | Spitz (SP) : Quantity Δ1 (1 unit) | 0.07 | 0.37 | 0.19 | 1185 | 0.85 | [-0.65 0.79] |
| | Golden Retriever (GR) : Quantity Δ1 (1 unit) | 0.18 | 0.17 | 1.07 | 1185 | 0.29 | [-0.15 0.52] |
| | Bichon Bolognese (BB): Quantity Δ1 (1 unit) | 0.03 | 0.15 | 0.20 | 1185 | 0.84 | [-0.27 0.33] |
| | Miniature Australian Shepherd (MAS) : Quantity Δ2 (2 units) | 0.21 | 0.12 | 1.67 | 1185 | 0.10 | [-0.04 0.45] |
| | White Swiss Shepherd (WS) : Quantity Δ2 (2 units) | 0.08 | 0.19 | 0.45 | 1185 | 0.66 | [-0.29 0.45] |
| | Jack Russell (JR) : Quantity Δ2 (2 units) | 0.10 | 0.29 | 0.33 | 1185 | 0.74 | [-0.48 0.67] |
| | Siberian Husky (SH) : Quantity Δ2 (2 units) | -0.07 | 0.15 | -0.46 | 1185 | 0.65 | [-0.37 0.23] |
| | Spitz (SP) : Quantity Δ2 (2 units) | -0.32 | 0.34 | -0.95 | 1185 | 0.34 | [-0.99 0.35] |
| | Golden Retriever (GR) : Quantity Δ2 (2 units) | -0.09 | 0.16 | -0.57 | 1185 | 0.57 | [-0.40 0.22] |
| | Bichon Bolognese (BB): Quantity Δ2 (2 units) | -0.01 | 0.15 | -0.07 | 1185 | 0.94 | [-0.30 0.28] |
| | (against French Bulldog (FB) and Quantity Δ3 (3 units)) | | | | | | |

*Supplementary table 2: Individual performance scores for the detection and discrimination tasks. The table contains proportion correct responses, the intercepts of the random factor subject and slopes grouped by side and task, and the ratio between the tasks:*

$$ratio = \frac{discrimination}{detection}$$

| Breed | Individual [#] | Detection | Discrimination | Ratio | ***Subject* x *Side*** | ***Subject* x *Task*** |
|---|---|---|---|---|---|---|
| Miniature Australian Shepherd (MAS) | 1 | 0.83 | 0.5 | 0.60 | | |
| | 2 | 0.67 | 0.22 | 0.33 | * | |
| | 3 | 0.78 | 0.39 | 0.50 | * | |
| | 4 | 0.89 | 0.61 | 0.69 | | |
| | 5 | 0.83 | 0.56 | 0.67 | | |
| | 6 | 0.44 | 0.33 | 0.75 | *** | |
| | 7 | 0.72 | 0.44 | 0.62 | *** | |
| | 8 | 0.94 | 0.50 | 0.53 | | |
| | 9 | 0.89 | 0.44 | 0.50 | | |
| White Swiss Shepherd (WS) | 1 | 0.83 | 0.61 | 0.73 | | |
| | 2 | 0.83 | 0.50 | 0.60 | | |
| | 3 | 0.89 | 0.50 | 0.56 | | |
| Jack Russell (JR) | 1 | 0.83 | 0.50 | 0.60 | | |
| | 2 | 1.00 | 0.67 | 0.67 | | |
| Siberian Husky (SH) | 1 | 1.00 | 0.33 | 0.33 | | |
| | 2 | 0.92 | 0.50 | 0.55 | | |
| | 3 | 0.83 | 0.58 | 0.70 | * | |
| | 4 | 0.75 | 0.67 | 0.89 | | |
| | 5 | 0.83 | 0.50 | 0.60 | | |
| | 6 | 0.92 | 0.42 | 0.45 | | |
| | 7 | 1.00 | 0.67 | 0.67 | | |
| | 8 | 0.92 | 0.67 | 0.73 | | |
| | 9 | 0.75 | 0.67 | 0.89 | | |
| Spitz (SP) | 1 | 0.75 | 0.50 | 0.67 | | |
| Golden Retriever (GR) | 1 | 0.78 | 0.44 | 0.57 | | |
| | 2 | 0.72 | 0.33 | 0.46 | | |
| | 3 | 0.61 | 0.44 | 0.73 | | |
| | 4 | 0.72 | 0.39 | 0.54 | | |
| | 5 | 0.78 | 0.61 | 0.79 | | |
| Bichon Bolognese (BB) | 1 | 0.33 | 0.33 | 1.00 | | |
| | 2 | 0.33 | 0.50 | 1.50 | * | |
| | 3 | 0.50 | 0.42 | 0.83 | * | |
| | 4 | 0.50 | 0.42 | 0.83 | *** | |
| | 5 | 0.17 | 0.42 | 2.50 | * | |
| | 6 | 0.33 | 0.42 | 1.25 | *** | |
| | 7 | 0.17 | 0.33 | 2.00 | | |
| | 8 | 0.83 | 0.17 | 2.00 | | |
| | 9 | 1.00 | 1.00 | 1.00 | | *** |
| French Bulldog (FB) | 1 | 0.94 | 0.42 | 0.44 | * | |
| | 2 | 0.67 | 0.92 | 1.38 | | |
| | 3 | 0.94 | 0.50 | 0.53 | * | |

Significance level [%]: * = 5, ** = 1, *** = .1

*Supplementary table 3: Results of the model investigating performance scores in detection and discrimination tasks. The table contains parameter estimates for the final model based on the factor 'cephalic index' (second model).*

| | | **Estimate** | **SE** | ***t*-stat** | **DF** | ***p*-value** | **CI (95%) [lower, upper]** |
|---|---|---|---|---|---|---|---|
| | Intercept | 0.34 | 0.11 | 3.22 | 1199 | 0.01 | [0.13 0.55] |
| **Cephalic index** | Cephalic index class 1 | -0.33 | 0.14 | -2.33 | 1199 | 0.02 | [-0.60 -0.05] |
| | Cephalic index class 2 | 0.06 | 0.17 | 0.33 | 1199 | 0.74 | [-0.28 0.39] |
| | (against Cephalic index class 3) | | | | | | |
| **Quantity** | Quantity Δ1 (1 unit) | -0.07 | 0.08 | -0.92 | 1199 | 0.36 | [-0.21 0.08] |
| | Quantity Δ2 (2 units) | -0.05 | 0.07 | -0.71 | 1199 | 0.47 | [-0.18 0.09] |
| | (against Quantity Δ3 (3 units)) | | | | | | |
| **Side** | Right | -0.15 | 0.10 | -1.61 | 1199 | 0.11 | [-0.34 0.03] |
| | (against Left) | | | | | | |
| **Task** | Detection task | 0.38 | 0.06 | 6.48 | 1199 | 0.001 | [0.26 0.49] |
| | (against Discrimination task) | | | | | | |
| **Cephalic index x Quantity** | Cephalic index class 1 : Quantity Δ1 (1 unit) | -0.04 | 0.11 | -0.34 | 1199 | 0.73 | [-0.24 0.17] |
| | Cephalic index class 1 : Quantity Δ2 (2 units) | -0.03 | 0.10 | -0.33 | 1199 | 0.74 | [-0.22 0.16] |
| | Cephalic index class 2 : Quantity Δ1 (1 unit) | 0.11 | 0.12 | 0.91 | 1199 | 0.36 | [-0.13 0.35] |
| | Cephalic index class 2 : Quantity Δ2 (2 units) | -0.07 | 0.11 | -0.69 | 1199 | 0.49 | [-0.29 0.14] |
| | (against Cephalic index class 3 and Quantity Δ3 (3 units)) | | | | | | |
| **Cephalic index x Task** | Cephalic index class 1 : Detection | -0.30 | 0.08 | -3.76 | 1199 | 0.001 | [-0.46 -0.15] |
| | Cephalic index class 2 : Detection | 0.12 | 0.09 | 1.31 | 1199 | 0.19 | [-0.06 0.31] |
| | (against Cephalic index class 3 and Discrimination task) | | | | | | |
| **Quantity x Task** | Quantity Δ1 (1 unit) : Detection | -0.05 | 0.07 | -0.69 | 1199 | 0.49 | [-0.17 0.08] |
| | Quantity Δ2 (2 units) : Detection | -0.01 | 0.07 | -0.10 | 1199 | 0.92 | [-0.14 0.12] |
| | (against Quantity Δ3 (3 units) and Discrimination task) | | | | | | |
